# Microbial paracetamol degradation involves a high diversity of novel amidase enzyme candidates

**DOI:** 10.1101/2022.05.05.490616

**Authors:** Ana B. Rios-Miguel, Garrett J. Smith, Geert Cremers, Theo van Alen, Mike S.M. Jetten, Huub J. M. Op den Camp, Cornelia U. Welte

## Abstract

Pharmaceuticals are relatively new to nature and often not completely removed in wastewater treatment plants (WWTPs). Consequently, these micropollutants end up in water bodies all around the world posing a great environmental risk. One exception to this recalcitrant conversion is paracetamol, whose full degradation has been linked to several microorganisms. However, the genes and corresponding proteins involved in microbial paracetamol degradation are still elusive. In order to improve our knowledge of the microbial paracetamol degradation pathway, we inoculated a bioreactor with sludge of a hospital WWTP (Pharmafilter, Delft, NL) and fed it with paracetamol as the sole carbon source. Paracetamol was fully degraded without any lag phase and the enriched microbial community was investigated by metagenomic and metatranscriptomic analyses, which demonstrated that the microbial community was very diverse. Dilution and plating on paracetamol-amended agar plates yielded two *Pseudomonas* sp. isolates: a fast-growing *Pseudomonas* sp. that degraded 200 mg/L of paracetamol in approximately 10 hours while excreting a dark brown component to the medium, and a slow-growing *Pseudomonas* sp. that degraded paracetamol without obvious intermediates in more than 90 days. Each *Pseudomonas* sp. contained a different highly-expressed amidase (31% identity to each other). These amidase genes were not detected in the bioreactor metagenome suggesting that other as-yet uncharacterized amidases may be responsible for the first biodegradation step of paracetamol. Uncharacterized deaminase genes and genes encoding dioxygenase enzymes involved in the catabolism of aromatic compounds and amino acids were the most likely candidates responsible for the degradation of paracetamol intermediates based on their high expression levels in the bioreactor metagenome and the *Pseudomonas* spp. genomes. Furthermore, cross-feeding between different community members might have occurred to efficiently degrade paracetamol and its intermediates in the bioreactor. This study increases our knowledge about the ongoing microbial evolution towards biodegradation of pharmaceuticals and points to a large diversity of (amidase) enzymes that are likely involved in paracetamol metabolism in WWTPs.

**Highlights:**

- Paracetamol was fully degraded by activated sludge from hospital wastewater.
- Low paracetamol concentrations were removed by a diverse microbial community.
- *Pseudomonas* sp. dominated cultures with high paracetamol concentration.
- Uncharacterized amidases are probably involved in degrading paracetamol in WWTPs.
- Deaminases and dioxygenases might be degrading paracetamol transformation products.

## 1. INTRODUCTION

Hundreds of pharmaceutical compounds are being detected at low concentrations in water bodies all around the world posing a severe risk to the environment and to human health (Gavrilescu et al., 2015; Wilkinson John et al., 2022). The consumption of medication and personal care products will most likely only increase in the future. Therefore, there is an urgent need to develop new technologies able to remove these chemicals at low concentrations before reaching the environment. Until now, cost-efficient removal of common pollutants (i.e. ammonium-nitrogen) has been achieved using microorganisms in wastewater treatment plants (WWTPs). As large-scale use of pharmaceuticals has only recently resulted in discharge to many different environments, the metabolic pathways of their conversion (at relatively low concentrations) might not have evolved yet or might not be very efficient.

Unlike many other pharmaceuticals, acetaminophen (N-acetyl-p-aminophenol, APAP), more commonly known as paracetamol, is degraded by microorganisms and is often fully removed in WWTPs. Several microorganisms have been related to APAP degradation in activated sludge and soil samples (i.e. *Penicillium, Pseudomonas, Flavobacterium, Dokdonella, Ensifer, Delftia*) (Hart and Orr, 1974; Palma et al., 2018; Park and Oh, 2020a; b; Rios-Miguel et al., 2021; Żur et al., 2018a). However, the genomes of these microorganisms have not yet been reported and therefore, the responsible genes and mechanisms for APAP biodegradation in WWTPs are not yet known.

4-Aminophenol (4-AP) and hydroquinone (HQ) have been measured in several APAP biodegradation experiments (Park and Oh, 2020a; b; Zhang et al., 2013). Consequently, an aryl acylamidase, a deaminase, and hydroquinone 1,2-dioxygenase were proposed as enzymes potentially involved in the biodegradation pathway of APAP (dos S. Grignet et al., 2022; Lee et al., 2015; Żur et al., 2018b). In fact, five amidases have been shown to transform APAP to 4-AP or to a brown compound (Supplementary Table S1) (Chen et al., 2016; Ko et al., 2010; Lee et al., 2015; Yun et al., 2017; Zhang et al., 2012; Zhang et al., 2020; Zhang et al., 2019). Besides, 1,2,4-trihydroxybenzene could be an intermediate of APAP degradation since it was measured in a *Burkholderia* sp. degrading 4-AP (Takenaka et al., 2003). Despite this knowledge, the exact genes/enzymes that microorganisms are using for APAP biodegradation in the environment are currently unknown.

To fill this gap, we analyzed the microbial community obtained from a hospital WWTP that degraded APAP in a bioreactor by metagenomics and metatranscriptomics. Furthermore, we were able to isolate two *Pseudomonas* species from the bioreactor that were capable of growing on APAP. Our aim was to identify the genes involved in APAP biodegradation and determine the genomic location and organization of these genes (clusters) in different microorganisms. These results will help to understand the evolution of microbial metabolism towards biodegradation of pharmaceuticals and will provide molecular biomarkers to screen environments for APAP-degrading microorganisms.

## 2. METHODS

### 2.1. Sampling and bioreactor set-up

Biomass was obtained from a membrane bioreactor (MBR) and a granular activated carbon (GAC) process at the Pharmafilter WWTP in Delft, the Netherlands, on 1-2-2021. This plant treats wastewater and solid waste from the Reinier de Graaf hospital, in Delft, and consists of an anaerobic-anoxic-oxic MBR, an ozonation tank, and a GAC treatment (https://www.stowa.nl/publicaties/evaluation-report-pharmafilter). A laboratory-scale membrane bioreactor (1.5 L) was inoculated with 15 mL of the MBR biomass and 15 mL of the GAC biomass. The lab-scale membrane and bioreactor vessel were built at Radboud University technical center. The bioreactor appliances were from Applikon Biotechnology B.V. (Delft, The Netherlands). The membrane consisted of an integral immersed Zenon ZW-1 module with 0.04 μm pore-sized hollow fibers from Suez Water Technologies & Solutions (Feasterville-Trevose, USA). It was never backwashed or replaced during the experiment. The bioreactor was fed with synthetic medium containing 0.05-0.4 g/L APAP (Merck, ≥99.0%, Darmstadt, Germany) as sole carbon source, 0.2 g/L K_2_HPO_4_, 0.1 g/L KH_2_PO_4_, 0.06 g/L NH4Cl, 0.01 g/L MgSO_4_ x 7 H_2_O, 0.01 g/L CaCl_2_, and trace elements solution (Rios-Miguel et al., 2021). Since APAP was degraded very fast, the organic loading rate was increased over the first 38 days from 0.022 to 0.227 mg APAP/min. This was done by increasing the concentration of APAP in the medium to 400 mg/L and by reducing the hydraulic retention time (HRT) from 2.4 to 1.8 d. When bacterial growth became exponential, the solid retention time (SRT) was set to 10 d to reach a steady state. After about 90 d, the HRT was set to 3.7 d to determine the microbial community changes at lower APAP loading rates. Furthermore, the bioreactor was run in the dark at constant 500 rpm stirring, a pH value of 7, an airflow rate of 30 ml/min, and room temperature (20 ± 1 °C). A pH sensor was connected to a controller that activated a KHCO3 base pump to keep the pH stable at 7.

### 2.2. Bioreactor monitoring: total suspended solids, paracetamol concentration, DNA, and RNA sequencing

Total suspended solids (TSS) were regularly measured by passing 30 mL of the sample through a 0.45 μm pore size glass-fiber filter which was dried overnight at 105 °C. Samples (2 mL) were taken regularly from the bioreactor in triplicates, centrifuged, and stored at −20 °C (both supernatant and pellet) until APAP and DNA analysis. APAP was measured in the supernatant using an HPLC-UV (Agilent Technologies 1000 series, injection volume of 100 μL; a mobile phase of acetate 1%: methanol (9:1); flow rate 1200 μl/min; and a C18 reverse-phase column: LiChrospher® 100 RP-18 (5 μm) LiChroCART® 125-4, 12.5 cm × 4 mm, Merck, Darmstadt, Germany). DNA was extracted from the pellets using the DNeasy PowerSoil Kit (Qiagen Benelux B.V.) following manufacturer’s instructions. The samples were submitted to Macrogen (Seoul, South Korea) for amplicon sequencing of the V3 and V4 regions of the bacterial 16S rRNA gene (primers Bac341F and Bac785R (Klindworth et al., 2013)) using an Illumina MiSeq. Six samples of 30 mL were taken at day 77 (two weeks after SRT was set to 10 d and the bioreactor was in a steady state). Three samples were centrifuged and stored at −20 °C for DNA sequencing and the other three were frozen in liquid nitrogen and stored at −80 °C for RNA sequencing. DNA was extracted using the DNeasy PowerSoil Kit (Qiagen Benelux B.V.) and sequenced at BaseClear (Leiden, The Netherlands). RNA was extracted using the RNeasy PowerSoil Total RNA Kit (Qiagen Benelux B.V.) with an extra DNase treatment from RibopureTM Kit (Thermo Fisher Scientific, Waltham, MA USA). Ribosomal RNA was removed using the Microbexpress kit (Life Technologies, Carlsbad, USA) and rRNA depleted samples were submitted to Macrogen (Seoul, South Korea) for sequencing. DNA and RNA samples were sequenced using Illumina Novaseq technology. All DNA and RNA quantities were determined using the Qubit dsDNA/RNA HS Assay Kit (Thermo Fisher Scientific, Waltham, MA USA) and a Qubit fluorometer (Thermo Fisher Scientific, Waltham, MA USA). Furthermore, DNA and RNA quality was checked with the Agilent 2100 Bioanalyzer and the High sensitivity DNA/RNA kit (Agilent, Santa Clara, USA).

### 2.3. Isolation and DNA/RNA sequencing of bacteria growing on paracetamol

We performed serial dilutions of the bioreactor biomass in synthetic medium (same as the bioreactor) with APAP as sole carbon source (0.2-0.4 g/L). The biomass of the most highly diluted culture displaying APAP biodegradation was plated on agar-solidified (1.5%) medium with APAP as sole carbon source. Two different colonies designed Pfast and Pslow were picked and inoculated in new bottles containing synthetic bioreactor medium (APAP sole carbon source). One milliliter of these cultures was used as inoculum for triplicate-bottle experiments where APAP biodegradation kinetics were measured and the DNA and RNA of the bacteria were sequenced. DNA and RNA from both isolates were extracted as described above, except for the Pfast DNA extraction for ONT sequencing, which was performed at Baseclear with the Wizard HMW DNA Extraction kit (Promega Benelux B.V., Leiden, The Netherlands). The genome of the fast-growing isolate Pfast was sequenced using Illumina Novaseq and ONT GridiON at Baseclear (Leiden, The Netherlands). RNA of this isolate was sequenced using Illumina Novaseq, also at Baseclear. Only one of the 3 RNA samples could be sequenced. The genome and transcriptome of the slow-growing isolate Pslow were sequenced using an in-house Illumina MiSeq. For DNA library preparation the Nextera XT kit was used and for transcriptomic library preparation, the TruSeq Stranded mRNA kit was used according to the manufacturer’s instructions (Illumina, San Diego, USA). All DNA and RNA quantifications and quality checks were performed as described above (Qubit and Bioanalyzer). For genomic DNA libraries, 300 bp paired-end sequencing was performed and for the transcriptomes, 150 bp single-read sequencing was done, using the Illumina Miseq sequencing machine (Illumina, San Diego, California). The raw sequence data and metadata of the bioreactor and isolates have been deposited at the read sequence archive (SRA) database of the NCBI under the BioProject ID PRJNA831879.

### 2.4. Bioinformatic analysis

#### 2.4.1. 16S rRNA gene sequencing data analysis

Analysis of the 16S rRNA sequencing output files was performed within R version 3.4.1 (Team, 2013) using the DADA2 pipeline (Callahan et al., 2016). Taxonomic assignment of the reads was up to the species level when possible using the Silva non-redundant database version 128 (Yilmaz et al., 2014). Count data were normalized to relative abundances. Data visualization and analysis were performed using phyloseq and ggplot packages (McMurdie and Holmes, 2013; Wickham and Wickham, 2007). Chao1, Simpson and Shannon diversity indices were calculated using the estimate richness function of the phyloseq package.

#### 2.4.2. DNA assembly, binning, and annotation

The quality of the metagenome sequencing data was assessed using FASTQC before and after quality processing. Quality-trimming, adapter removal and contaminant filtering of Illumina paired-end sequencing reads was performed using BBDuk (BBTools, DOE Joint Genome Institute, Lawrence Berkeley National Laboratory, USA). The DNA trimmed reads were assembled using MetaSpades and aligned to the assembly using BBMap to generate coverage information (Nurk et al., 2017). The assemblies were binned using different binning algorithms (BinSanity, CONCOCT, MaxBin 2.0, and MetaBAT 2) (Alneberg et al., 2014; Graham et al., 2017; Kang et al., 2019; Wu et al., 2015). DAS Tool was used for consensus binning (Sieber et al., 2018). GTDB-Tk was used to assign taxonomy and CheckM was used to assess the quality of the bins or metagenome-assembled genomes (MAGs) (Chaumeil et al., 2019; Parks et al., 2015). Annotation was performed by Metascan (Cremers et al. under revision). The annotation method was described previously by ‘t Zandt et al. and Poghosyan et al. (in ’t Zandt et al., 2020; Poghosyan et al., 2020). Briefly, genes were called by Prodigal (Hyatt et al., 2010) and subsequent open reading frames were annotated with HMMER (Eddy, 2009) and a set of custom-made databases.

DNA sequencing data from the slow-growing isolate Pslow were quality-checked and trimmed with BBDuk, assembled with MetaSpades and annotated with Metascan. The DNA sample consisted of two genomes (a very minor contaminant) so we separated the contigs using Maxbin 2.0. (Wu et al., 2015).

DNA Illumina reads from the fast-growing isolate Pfast were quality controlled and trimmed using BBduk with minimum trim quality of 18 and length of 100. ONT reads were filtered to a minimum length of 3000 using BBtools utilities, and then were quality controlled and trimmed using Porechop (https://github.com/rrwick/Porechop) with minimum split read size of 3000. Illumina and ONT quality-trimmed reads were assembled using Unicycler with a minimum length of 1000 in the resulting. Metascan was used for annotation. Completion and contamination scores of the Pslow and Pfast assemblies were estimated using CheckM’s lineage workflow. The whole-genome phylogenetic position of both assemblies was inferred using GTDB-tk. If not specified, settings were default.

#### 2.4.3. Transcriptome analysis

The RNA reads were trimmed using Sickle and mapped with BBMap (allowing 1% mismatch, BBtools) to the protein-coding genes of each isolate or to the contigs of the whole metagenome. Then, transcripts per million (TPM) were calculated in Excel for each gene in each sample: first, reads per kilobase were calculated (read counts divided by the length of each gene in kilobases); second, the “per million” scaling factor was calculated (sum of all the reads per kilobase in one sample divided by one million); and third, TPM were calculated (reads per kilobase of each gene divided by the “per million” scaling factor). Approximately, the top 10% most highly expressed genes in each microorganism or bin were considered as “highly-expressed”. The RNA coverage of each bin in the metagenome was calculated or defined as the number of bases mapped to the set of protein-coding genes from each bin divided by the total number of bases in each bin. Amidase sequences were retrieved by searching for “amidase”, “amidohydrolase”, and “amidotransferase” terms in the annotated metagenome. The highly-expressed and uncharacterized amidase sequences were aligned with Clustal Omega (Higgins and Sharp, 1988) and MUSCLE (Edgar, 2004) and the phylogenetic trees were created with MEGA7 (Kumar et al., 2016) and MEGA11 (Tamura et al., 2021) using the maximum likelihood algorithm (Jones et al., 1992) and bootstrap analysis or the neighbor-joining method to analyse the tree topology (Saitou and Nei, 1987).

## 3. RESULTS AND DISCUSSION

### 3.1. Bioreactor performance and bacterial community changes

A bioreactor was inoculated with sludge from a WWTP treating hospital waste. APAP was added as the sole carbon source at all times during the experiment and it was fully degraded since the beginning without lag phase. No transformation products were detected. The microbial community was very diverse and changed during the different operational settings (Figure 1). In the first start-up phase, the total suspended solids (TSS) gradually increased to about 1.1 g/L as all biomass was retained in the bioreactor via a membrane module. The HRT was 1.8 days. Under these conditions, the microbial community was dominated by members of the families *Chitinophagaceae*, *Haliangiaceae, Phycisphaeraceae, Pseudomonadaceae, Saccharimonadaceae*, and *Sphingobacteriaceae* when compared to the inoculum mix. In the second phase, a SRT of 10 days was maintained in order to keep the biomass concentration (TSS) stable at a steady state. This led to a decrease of the relative abundance of *Pseudomonadaceae*, *Phycisphaeraceae*, and *Sphingobacteriaceae* while increasing *Burkholderiaceae*, *Chitinophagaceae*, and *Polyangiaceae*. In the third phase, the HRT was increased from 1.8 to 3.7 days resulting in a bacterial community dominated by the heterotrophic bacteria *Blastocatellaceae*, *Burkholderiaceae*, and *Chitinophagaceae*. *Nitrospiraceae* were also enriched which might be the result of growth on low concentration of residual ammonium in the reactor. Overall, the alpha diversity (richness and evenness of species) decreased over time in the bioreactor which corresponds to a selection and enrichment process (Figure S1). The previously mentioned taxa are normally found in WWTPs due to their ability to degrade organic matter or ammonium/nitrite (*Nitrospira*) (Morin et al., 2020; Saunders et al., 2016). The presence of heterotrophs able to degrade complex organic matter (i.e. *Chitinophagaceae* and *Polyangiaceae*) might indicate possible predation and biomass recycling in the bioreactor (Petters et al., 2021). Furthermore, members of the *Pseudomonadaceae* and *Burkholderiaceae* families have been reported to degrade APAP and 4-AP, respectively (Park and Oh, 2020a; Takenaka et al., 2003; Żur et al., 2018a).

**Figure 1.**
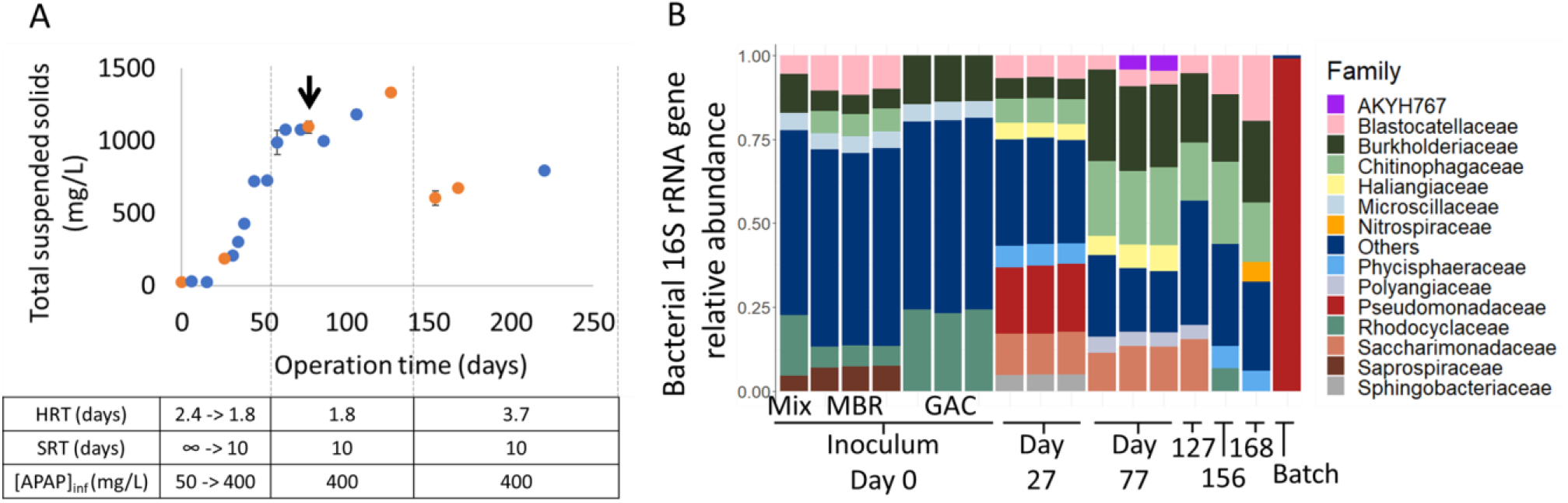
**A**: Total suspended solids in mg/L. Orange dots represent the time points when 16S rRNA genes were sequenced. The black arrow represents the time point when whole-genome and transcriptome sequencing occurred. **B**: relative abundance of bacterial 16S rRNA genes in the inoculum and the bioreactor at several time points. Batch sample represents the serial dilution of the bioreactor biomass in medium containing 400 mg/L of APAP. Abbreviations: HRT, hydraulic retention time; SRT, solid retention time; APAP, paracetamol; MBR, membrane bioreactor; GAC, granular activated carbon; Mix, mixture of MBR and GAC.

### 3.2. Recovery of metagenome-assembled genomes from the bioreactor

Fourteen MAGs were recovered from the bioreactor metagenome and approximately 30% of the total reads remained unbinned. Table 1 shows the 14 recovered MAGs ordered from highest to lowest coverage based on RNA sequencing data (calculated with RNA bases mapped to protein-coding genes). *Chitinophagaceae* and *Myxococcales* were the most active (RNA coverage) bacteria and also the most abundant (DNA coverage) together with *Microbacterium* and *Patescibacteria*. Despite the high abundance of the *Patescibacteria* MAG, it had low completeness. The reason for this might be that the single copy genes normally used to calculate completeness are often not detected in *Patescibacteria* genomes (Brown et al., 2015). *Pseudomonas* spp. were low abundant in the metagenome and only present in the unbinned reads.

**Table 1.**
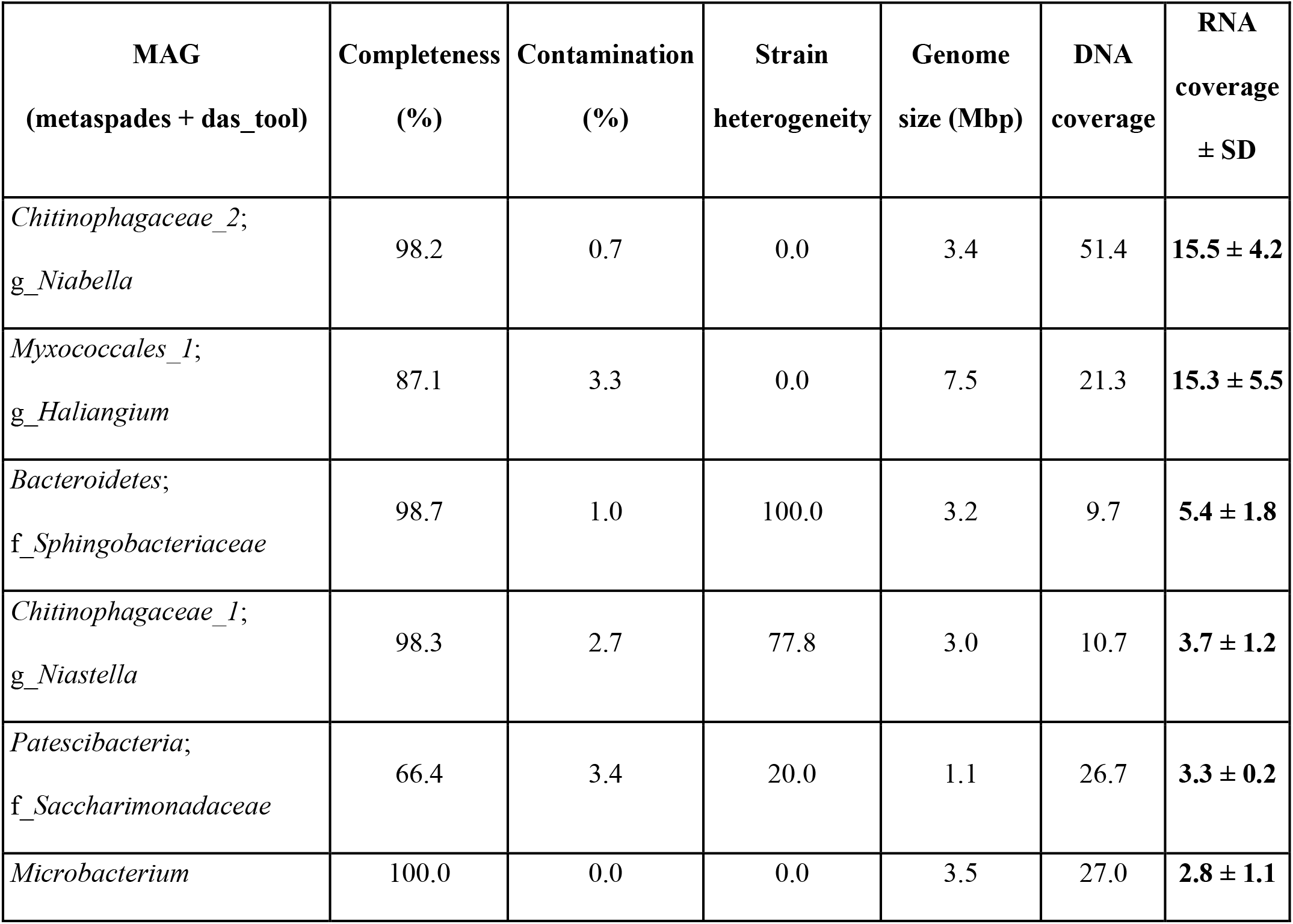
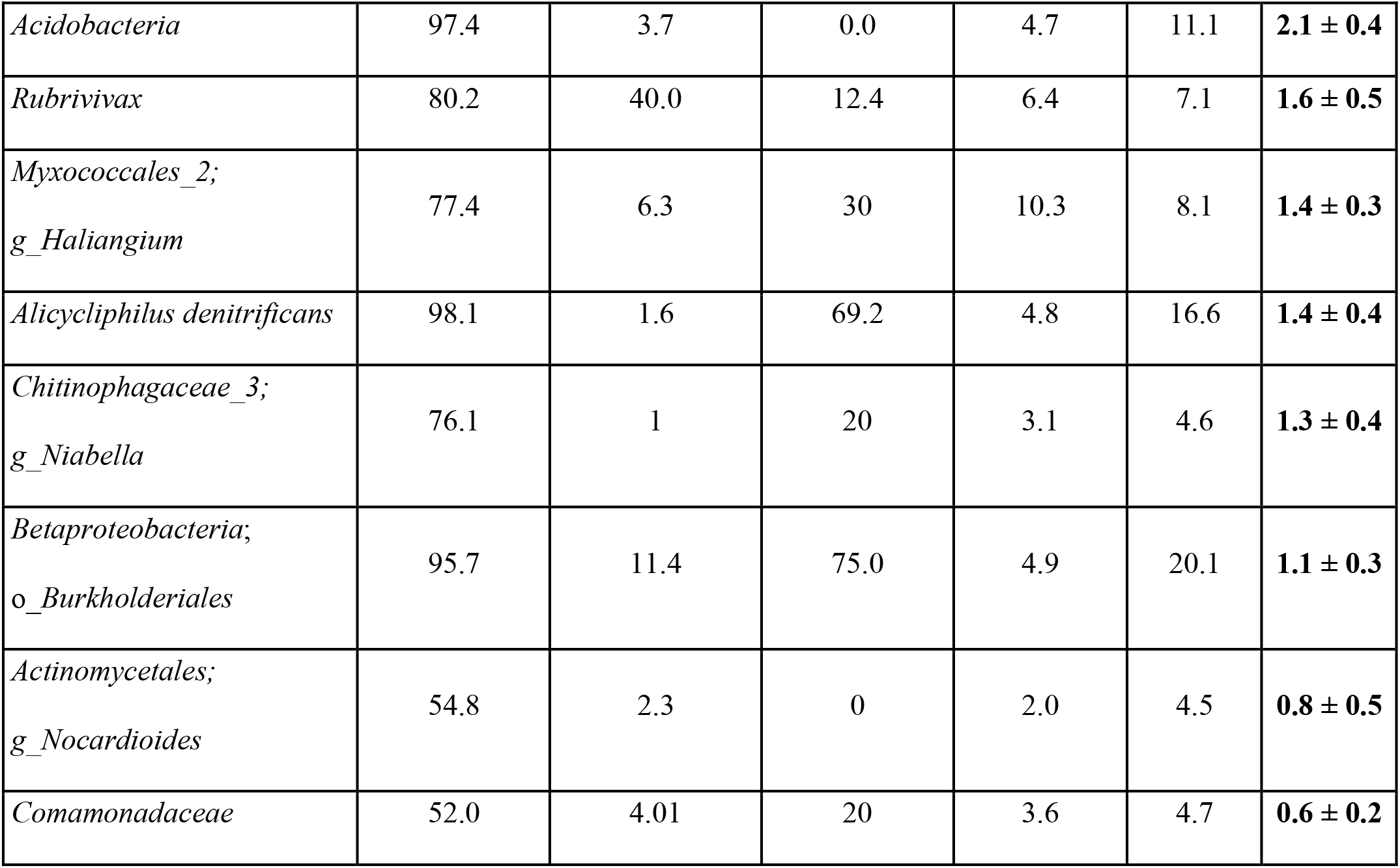
Metagenome-assembled genomes (MAGs) from the bioreactor at day 77. CheckM was used to check the quality of the MAGs. Taxonomy was assigned until the highest level possible using GTDB-Tk. RNA coverage is the average of three replicate samples while DNA coverage is based on one sample.

### 3.3. Paracetamol degradation by two *Pseudomonas* spp. isolates

After serial dilutions of the bioreactor biomass, and plating the highest dilution showing growth on agar mineral medium with APAP as sole carbon source, two *Pseudomonas* spp. were obtained (Figure 1B (Batch) and Figure 2). *Pseudomonas* sp. Pfast degraded 200 mg/L APAP in 10 h and converted it into 4-AP, which precipitated in the medium as a dark-brown solid. 4-AP might also be degraded by the Pfast isolate much slower than APAP. HQ was also detected as an intermediate but in a much lower concentration than 4-AP (data not shown). The other *Pseudomonas* sp., Pslow, degraded 200 mg/L APAP in approximately 90 d without the accumulation of aromatic transformation products. Previous studies showed that other *Pseudomonas* spp. are also able to degrade APAP in a few hours (De Gusseme et al., 2011; Hu et al., 2013; Park and Oh, 2020a; Zhang et al., 2013; Żur et al., 2018a). However, we are not aware of reports describing bacteria that degrade APAP at low rates.

**Figure 2.**
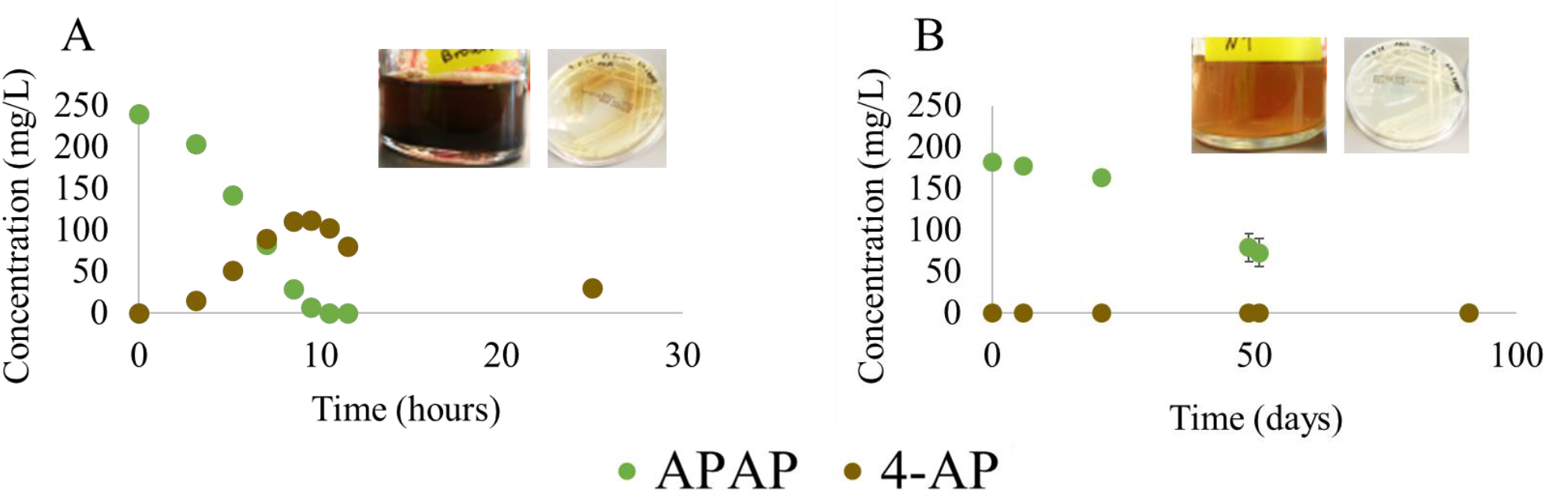
APAP biodegradation rates of two Pseudomonas spp. isolated from the bioreactor by serial dilutions and plating. **A** corresponds to the fast-growing Pseudomonas sp. Pfast and **B** to the slow-growing Pseudomonas sp. Pslow. Abbreviations: APAP, paracetamol; 4-AP, 4-aminophenol.

### 3.4. Highly-expressed amidases in the two *Pseudomonas* spp. isolates

After DNA and RNA sequencing, a highly expressed gene cluster was identified in the Pfast isolate that contained a putative amide transporter (AmiS/UreI family, OACKLNDA_05759) and an amidase-like protein (OACKLNDA_05760) with 85% sequence identity to an aryl acylamidase known to convert APAP into 4-AP and acetate (Ko et al., 2010; Lee et al., 2015). The transporter might be catalyzing the uptake of APAP or the excretion of 4-AP. The genome of the Pslow isolate contained neither this amidase nor the putative amide transporter. Instead, it encoded three other amidases with ≥ 95% query coverage and 28-31% identity to the Psfast amidase. The most similar amidase (31% identity, BKDBLJDL_05334) was upregulated in the transcriptome of strain Pslow cultivated with APAP as the sole carbon source. This amidase was encoded in a gene cluster together with a gene for a zinc-dependent hydrolase (BKDBLJDL_05335). Interestingly, these genes were also present in the fast-growing *Pseudomonas* isolate, but not highly expressed. The difference in APAP biodegradation rate and consequently, growth between both Pseudomonas spp. was most likely related to the presence of OACKLNDA_05760 amidase in Pfast, which might be able to transform APAP to 4-AP at high rates. Another option could be that the putative amide transporter OACKLNDA_05759 was involved in a faster uptake of APAP and thus, transformation into 4-AP. Further studies are needed to answer this question.

The highly expressed amidase gene of strain Pslow was present in the chromosome of numerous *Pseudomonas* species registered in the NCBI nucleotide collection. However, the highly expressed amidase gene from strain Pfast was only found in a few microorganisms that came from various locations around the world (Australia, China, Pakistan, India, Korea, India, Poland): in the plasmid of multi-drug resistant *Acinetobacter* spp. isolated from patients and hospitals (Ghaly et al., 2020; Kizny Gordon et al., 2020; Zou et al., 2017); and in the chromosome of *Pseudomonas* and *Burkholderia* spp. isolated from soil, activated sludge, and hospitals (D’Souza et al., 2019; Ko et al., 2010; Patil et al., 2017; Żur et al., 2018b). Many bacteria contained a similar AmiS/UreI family transporter gene next to the amidase gene and different mobile genetic elements nearby (Tn3 transposons, *IS630* insertion sequences and IntI1 integrases). Our strain, Pfast, also had two insertion sequences *(IS6100* and *IS21*, OACKLNDA_05756, OACKLNDA_05781), one Tn3 transposase gene (OACKLNDA_05752), and one recombinase gene (OACKLNDA_05755) near the highly-expressed amidase gene. Consequently, the whole gene cluster might have been exchanged between different species via horizontal gene transfer (HGT) (Rios Miguel et al., 2020). Finally, the low number of homologous proteins in the NCBI database might indicate the recent evolution of this amidase towards paracetamol biodegradation or our limited ability to find and identify these genes.

### 3.5. Amidase diversity in the metagenome

To check and confirm whether the metagenome contained more unidentified amidases involved in APAP and intermediate conversion, we analyzed the total metagenome in more detail. The highest BLAST identity match of the Pfast amidase was 50% for proteins encoded on the unbinned contigs. The 14 MAGs only contained amidases with a maximum of 30% sequence identity. For the Pslow isolate, the highest match was 50% for proteins encoded on the unbinned contigs as well as the *Rubrivivax* and *Betaproteobacteria* MAGs. Since the APAP concentration was below the detection limit (~0.2 mg/L) in the bioreactor, but continuously supplied with the medium inflow, uncharacterized amidases might be responsible for APAP biodegradation at low concentrations in the bioreactor.

A phylogenetic tree was created with the top 150 most expressed amidases in the metagenome, the uncharacterized amidases (enzymes annotated as “amidase” and whose function is not known) present in both *Pseudomonas* isolates, and five amidases known to degrade APAP obtained from literature and databases (Supplementary Figure S2, Supplementary Table S1). All the uncharacterized amidases from the metagenome and the *Pseudomonas* isolate genomes clustered together (green cluster in Supplementary Figure S2), including four amidases known to degrade paracetamol (Ko et al., 2010; Lee et al., 2015; Yun et al., 2017; Zhang et al., 2012; Zhang et al., 2020; Zhang et al., 2019). This group belongs to the Amidase Signature (AS) enzyme family [EC:3.5.1.4] characterized by a highly conserved signature region of approximately 160 amino acids that includes a canonical catalytic triad (Ser-*cis*Ser-Lys) and a Gly/Ser-rich motif (GGSS[GS]G). One amidase (AGC74206.1 dimethoate hydrolase DmhA), previously reported to degrade APAP (Chen et al., 2016), did not cluster in this group (Supplementary Figure S2). Consequently, there is the possibility that other amidase families are also able to transform APAP. For instance, a histone deacetylase-like amidohydrolase clustered together with the APAP-degrading amidase DmhA, suggesting its reactivity towards APAP.

The green AS amidase cluster of the phylogenetic tree in Supplementary Figure S2 was analyzed in more detail (Figure 3). The amidase gene expression level in each bin/MAG was added to the tree (i.e. top 10%) and the amino acid sequences were manually blasted against the NCBI non-redundant protein database to improve the annotation. Furthermore, partial sequences like one highly-expressed amidase gene from *Betaproteobacteria* (EAEDEFLN_03924) were removed from the analysis. Twelve amidases were identified as “Asp-tRNA(Asn)/Glu-tRNA(Gln) amidotransferase subunit GatA” and they all clustered together (green cluster in Figure 3). This type of amidase is involved in the transformation of Glu-tRNA^Gln^ to Gln-tRNA^Gln^ for the synthesis of proteins. Gene duplication and mutation events in this amidotransferase were probably the key contributors to the high number of uncharacterized amidases with broad substrate specificity.

Another cluster, with low bootstrap support, in the phylogenetic tree of Figure 3 contained highly-expressed amidases (top 10%) of the *Pseudomonas* genomes and the *Microbacterium* MAG (red amidases in Figure 3, OACKLNDA_05760, BKDBLJDL_05334, BJLACCKG_02946). The *Microbacterium* amidase only had 91% query coverage and 63% identity to the closest amidase in the NCBI non-redundant protein database. This means that this amidase was sequenced for the first time and thus, the evolution of amidases towards paracetamol biodegradation might be an ongoing process. The *Microbacterium* amidase was part of a highly-expressed gene cluster containing a flavin reductase (BJLACCKG_02944), an arylformamidase (BJLACCKG_02945), a branched-chain amino acid ABC transporter (BJLACCKG_02947, BJLACCKG_02948, BJLACCKG_02949, BJLACCKG_02950, BJLACCKG_02951), and one oxidoreductase (BJLACCKG_02952). Furthermore, an *IS*110 family transposase gene (BJLACCKG_02943) was right next to this highly expressed gene cluster. Four amidases known to degrade APAP were also part of the phylogenetic tree cluster containing the highly-expressed amidases of the *Pseudomonas* genomes and *Microbacterium* MAG (ANS81375.1 arylamidase Mah, ACP39716.2 aryl acylamidase, ANB41810.1 tricocarban amidase TccA, AFC37599.1 aryl-amidase A AmpA; blue amidases in Figure 3). Three amidase genes present on the unbinned contigs were highly expressed in relation to all the unbinned protein-coding genes (top 10%, orange amidases in Figure 3). They were affiliated with *Actinomycetia* and *Comamonadaceae* spp. (KGEGGEOM_07080, KGEGGEOM_08489, KGEGGEOM_08132) and might also be degrading APAP in the bioreactor. Non-highly-expressed amidase genes might also have the potential to degrade APAP even though their genes were not strongly regulated when APAP was present. For instance, the non-highly-expressed amidase OACKLNDA_00714 of the Pfast strain was identical to the highly-expressed amidase BKDBLJDL_05334 of the Pslow strain, so it had the potential to degrade APAP at slow rates but it was not highly-expressed in Pfast.

Finally, a multiple sequence alignment was performed with the amidases known to degrade APAP (except for DmhA which is not part of the AS family) and the highly expressed amidases from the *Pseudomonas* genomes and the metagenome (Supplementary File 2). Lee et al. previously determined the three-dimensional structure of the amidase ACP39716.2 with APAP as a substrate (Lee et al., 2015). They revealed several residues involved in catalysis and APAP binding that we investigated in our alignment. The aligned amidases contained a conserved catalytic triad (Ser^187^-*cis*Ser^163^-Lys^84^, highlighted in green), Gly/Serrich motif (GGSSGG, in bold) and oxyanion hole ([G]GGS, in bold). The substrate-binding pocket contained two loop regions (highlighted in fair and dark grey) and one α-helix (highlighted in blue) that were less conserved. In the crystal structure of ACP39716, another two residues (Tyr^136^ and Thr^330^, highlighted in yellow) were described to bind to the hydroxyl group at the para-position in APAP via hydrogen bonds with two water molecules. However, Thr^330^ was only present in approximately half of the aligned amidase sequences and Tyr^136^ was not present in any of them. Thus, the amidases from this study might have different substrate specificities compared to the ACP39716.2 amidase. Furthermore, we conclude that Tyr^136^ and Thr^330^ are not strictly necessary for APAP binding and degradation.

**Figure 3.**
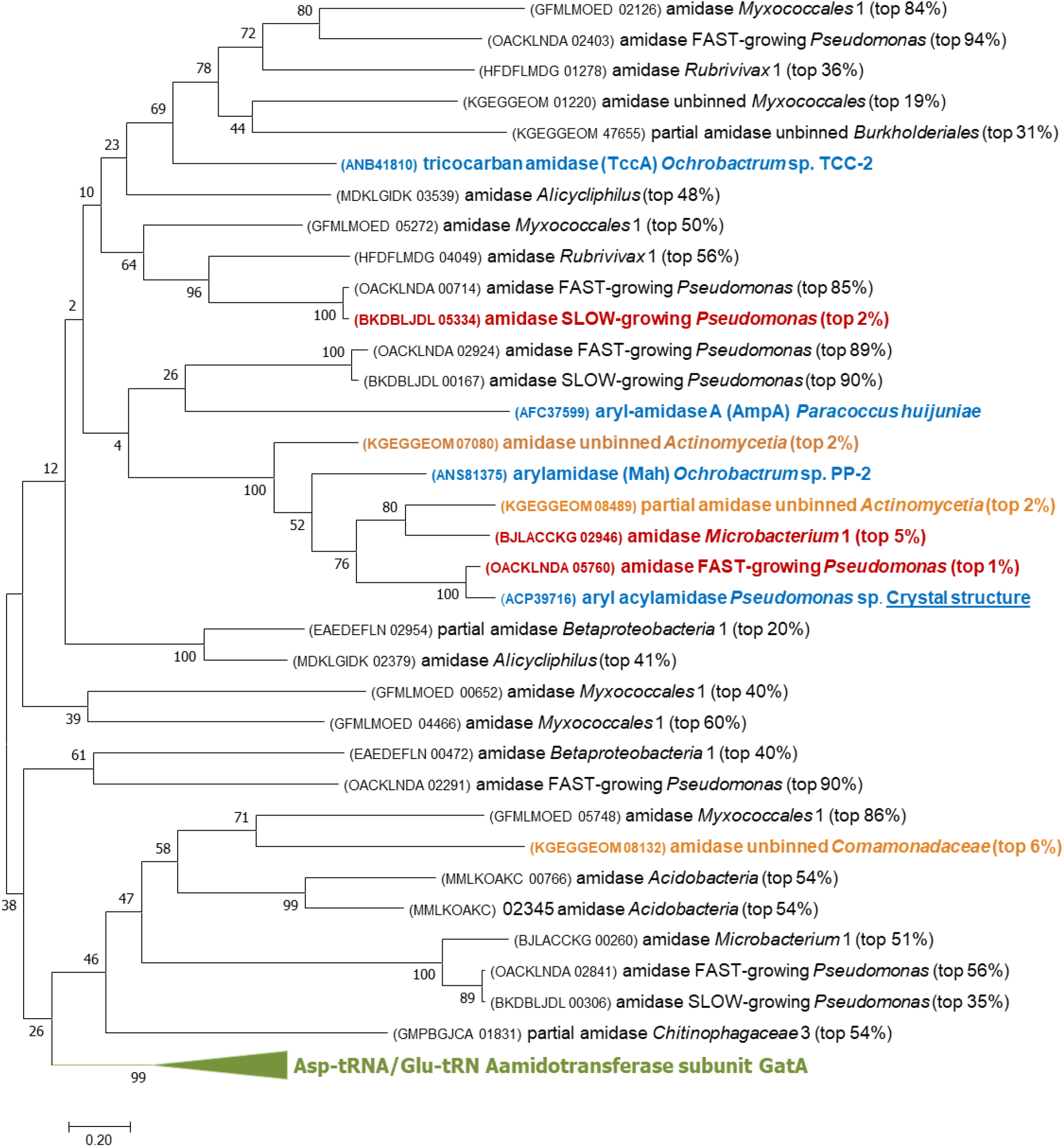
Phylogenetic tree of the Amidase Signature (AS) enzyme family [EC:3.5.1.4] proteins in the bioreactor and the Pseudomonas isolates. The evolutionary history was inferred by using the Maximum Likelihood method (Jones et al., 1992). The tree with the highest log likelihood (−25739.55) is shown after bootstrapping 500 times. The percentage of trees in which the associated taxa clustered together is shown next to the branches. Evolutionary analyses were conducted in MEGA7 (Kumar et al., 2016). Amidases in blue are experimentally validated to degrade APAP. Amidases in red are the ones whose expression lies in the top 10% of all the genes in the Pseudomonas isolate transcriptomes, and the metatranscriptomes of the bioreactor mapping to a metagenome-assembled genome in this study. Orange amidases correspond to the amidase genes lying in the top 10% most expressed from the unbinned protein-coding genes. The green cluster corresponds to amidases annotated as the Asp-tRNA(Asn)/Glu-tRNA(Gln) amidotransferase subunit GatA.

### 3.6. Paracetamol-degradation pathway: highly-expressed gene candidates

The first step in APAP biodegradation is the cleavage of the amide bond by an amidase to produce 4-AP and acetate (**Figure 4**). In each *Pseudomonas* genome, a different highly expressed amidase was identified presumably performing this cleavage (OACKLNDA_05760, BKDBLJDL_05334). In the bioreactor metagenome, the *Microbacterium* MAG was the only one with a highly expressed amidase inside the AS family cluster (BJLACCKG_02946), from which four amidases were previously reported to degrade APAP (**Figure 3**). The *Betaproteobacteria* MAG also had a highly expressed uncharacterized amidase gene (EAEDEFLN_03924). However, the nucleotide sequence was partial so we could not check its classification. Therefore, *Microbacterium* (and *Betaproteobacteria*) might have been involved in transforming APAP into 4-AP together with some low abundant bacteria in the unbinned group.

The enzyme deaminating 4-AP is still unknown and we did not find an obvious gene responsible for this reaction. Uncharacterized RidA family protein genes were highly expressed in the two *Pseudomonas* genomes and in the *Microbacterium* and *Betaproteobacteria* MAGs (OACKLNDA_05815, OACKLNDA_03065, BKDBLJDL_03266, BKDBLJDL_01320, BJLACCKG_02940, BJLACCKG_02936, EAEDEFLN_02051). Therefore, these proteins might have been involved in deaminating 4-AP or deaminating the aminomuconate intermediates after ring cleavage (He and Spain, 1998). Furthermore, two ammonia-lyase genes were highly expressed in the genome of the Pslow strain: aspartate ammonia-lyase and ethanolamine ammonia-lyase (BKDBLJDL_01355, BKDBLJDL_02838). However, these genes were not highly expressed in the Pfast genome and the metagenome from the bioreactor.

The investigated bacteria did not use any known hydroquinone 1,2-dioxygenase, which is a type III ring-cleaving extradiol dioxygenase (cupin superfamily) with a catalytic mechanism analogous to that of the extradiol-type dioxygenases. Instead, the up-regulated type III extradiol dioxygenase genes present in bioreactor MAGs (*Chitinophagaceae*_1,2 and *Betaproteobacteria*: 3-hydroxyanthranilate 3,4-dioxygenase (NLCKOFOF_01117, JPNLMFAJ_02227, EAEDEFLN_00964); *Bacteroidetes*, *Rubrivivax*, and *Myxococcales*_2: homogentisate 1,2-dioxygenase (KGEBAGMN_02738, HFDFLMDG_05905, OPIJCCOK_08966)) might have cleaved the aromatic ring of HQ to produce 4-hydroxymuconic semialdehyde as previously described (Żur et al., 2018b) (**Figure 4**). Homogentisate 1,2-dioxygenase and 3-hydroxyanthranilate 3,4-dioxygenase are type III extradiol dioxygenases able to cleave the aromatic ring of non-catecholic substrates, which are characterized by not having vicinal diols. For example, (homo)gentisate and hydroquinone have two hydroxyl groups in *para* position and 3-hydroxyanthranilate has one hydroxyl, one amino, and one carboxylic acid group as ring substituents. These two dioxygenases are involved in the degradation of aromatic amino acids. Therefore, bacteria might use the side activities of existing enzymes to degrade aromatic micropollutants such as APAP.

The ring cleavage of aromatic compounds containing hydroxyl and amino substituents (i.e. 2-aminophenol, 3-hydroxyanthranilate, and 5-aminosalicylate) has been previously reported (Hintner et al., 2004; Li de et al., 2013; Takenaka et al., 1997; Wang et al., 2020). Therefore, the aromatic ring of 4-AP might also be cleaved before deamination by class III ring-cleaving dioxygenases. However, metabolites confirming this pathway have not yet been measured.

An alternative route for the direct ring cleavage of HQ is the hydroxylation of HQ to form hydroxyquinol (1,2,4-trihydroxybenzene) and later, the ring cleavage of hydroxyquinol by an intradiol-type dioxygenase (Ferraroni et al., 2005; Takenaka et al., 2003) or an extradiol-type dioxygenase, probably from the vicinal oxygen chelate (VOC) or type I superfamily (Murakami et al., 1999) (**Figure 4**). The hydroxylation of HQ can be performed by ring-hydroxylating dioxygenases or monooxygenases. In the Pfast isolate genome, an uncharacterized ring-hydroxylating dioxygenase (OACKLNDA_03401) and an intradiol catechol 1,2-dioxygenase (OACKLNDA_05722) were highly expressed. Furthermore, two uncharacterized extradiol-type dioxygenases were highly expressed in both *Pseudomonas* isolate genomes (OACKLNDA_01459, BKDBLJDL_01807). These dioxygenases were similar to 4,5-DOPA dioxygenase, which is part of the VOC extradiol dioxygenases (Wang et al., 2019). In the bioreactor, the *Rubrivivax* MAG had a phenol hydroxylase (HFDFLMDG_03468, HFDFLMDG_03467, HFDFLMDG_03466), an uncharacterized ring-hydroxylating dioxygenase (HFDFLMDG_03024), a 4-hydroxybenzoate 3-monooxygenase (HFDFLMDG_05635), and a extradiol protocatechuate 4,5-dioxygenase (HFDFLMDG_01086, HFDFLMDG_01087) highly expressed. The *Microbacterium* MAG contained a highly-expressed VOC extradiol 3,4-dihydroxyphenylacetate (homoprotocatechuate) 2,3-dioxygenase involved in the degradation of tyrosine (BJLACCKG_03009). The *Betaproteobacteria* MAG had a putative ring hydroxylating dioxygenase (EAEDEFLN_00364), and a 4-hydroxyphenylpyruvate dioxygenase (EAEDEFLN_02889) able to hydroxylate and decarboxylate aromatic rings in the tyrosine degradation pathway. Many other MAGs contained this gene up-regulated, i.e. *Bacteroidetes*, *Comamonadaceae* and *Myxococcales*_2 (KGEBAGMN_02075, JMGBCLMB_02041, OPIJCCOK_08967). Finally, the *Alicycliphilus* MAG had a highly expressed gene cluster containing an MFS transporter (MDKLGIDK_02282), a tripartite tricarboxylate transporter substrate binding protein (MDKLGIDK_02283), an muconolactone D-isomerase (MDKLGIDK_02284), an 3-oxoadipate enol-lactonase (MDKLGIDK_02285), a 1,6-dihydroxycyclohexa-2,4-diene-1-carboxylate dehydrogenase (MDKLGIDK_02286), an intradiol catechol 1,2-dioxygenase gene (MDKLGIDK_02287), and a muconate cycloisomerase (MDKLGIDK_02288). This highly-expressed gene cluster suggests the ability of *Alicycliphilus* to fully degrade hydroxyquinol via the oxoadipate pathway. Interestingly, the *Alicycliphilus* MAG had hydroquinone dioxygenase genes (MDKLGIDK_00653, MDKLGIDK_00654) that were not highly expressed.

Acetyl-CoA synthetase genes and genes encoding enzymes involved in the tricarboxylic acid (TCA) and glyoxylate cycles (isocitrate lyase) were highly expressed in the transcriptome of both *Pseudomonas* spp. and several MAGs (i.e. *Microbacterium*, *Rubrivivax*, *Alycicliphilus*, and *Actinomycetales*), thus indicating their ability to grow on acetate after cleaving the APAP amide bond or via cross-feeding from other bacteria (**Figure 4**). Finally, the highly-expressed gene encoding an uncharacterized carboxymuconolactone decarboxylase family protein could be involved in the conversion of muconolactone intermediates to eventually reach the TCA cycle for bacterial growth in the *Pseudomonas* isolates (OACKLNDA_03472, BKDBLJDL_01962). The *Microbacterium*, *Rubrivivax*, and *Betaproteobacteria* MAGs had highly-expressed dioxygenases but they did not have up-regulated genes belonging to the β-ketoadipate pathway, involved in the conversion of aromatic metabolites into TCA intermediates. Therefore, it is unclear whether they were able to assimilate aromatic compounds or not. Furthermore, the *Rubrivivax* and *Alicycliphilus* MAGs did not have any highly-expressed amidase from the AS family suggesting that cross-feeding of acetate and aromatic intermediates was happening in the bioreactor.

The *Myxococcales* family is known for its diverse metabolism and its predatory nature, so the two bioreactor MAGs affiliated to the *Myxococcales* family might have lived from the metabolites and cellular components of decaying microorganisms (Müller et al., 2016). Similarly, the *Chitinophagaceae* family is known to degrade complex organic matter and therefore, the three bioreactor MAGs affiliated to the *Chitinophagaceae* family could also have been biomass recyclers in the bioreactor (Morin et al., 2020). The highest expressed metabolic genes of these MAGs were involved in the TCA cycle, gluconeogenesis, and metabolism of lipids, peptidoglycan, nucleotides, and amino acids, thus not providing many hints about their exact catabolism or energy source. Similarly, the metabolism of *Bacteroidetes, Acidobacteria, Actinomycetales*, and *Comamonadaceae* MAGs was ambiguous and they might also be predators or biomass recyclers. In addition, we found that type II and type IV secretion systems were highly expressed in several MAGs (i.e. *Acidobacteria, Comamonadaceae*) which might have been involved in predation, defense, and conjugation activities between microorganisms (Aharon et al., 2021; Sgro et al., 2019). However, some of these MAGs might also degrade APAP transformation products via their highly-expressed dioxygenases (i.e. *Chitinophagaceae*_1_2, *Bacteroidetes*, *Myxococcales*_2). The *Patescibacteria* MAG mostly contained genes encoding carbohydrate degrading enzymes, so it might have thrived in symbiosis with other microbial community members that produced exopolysaccharides.

**Figure 4.**
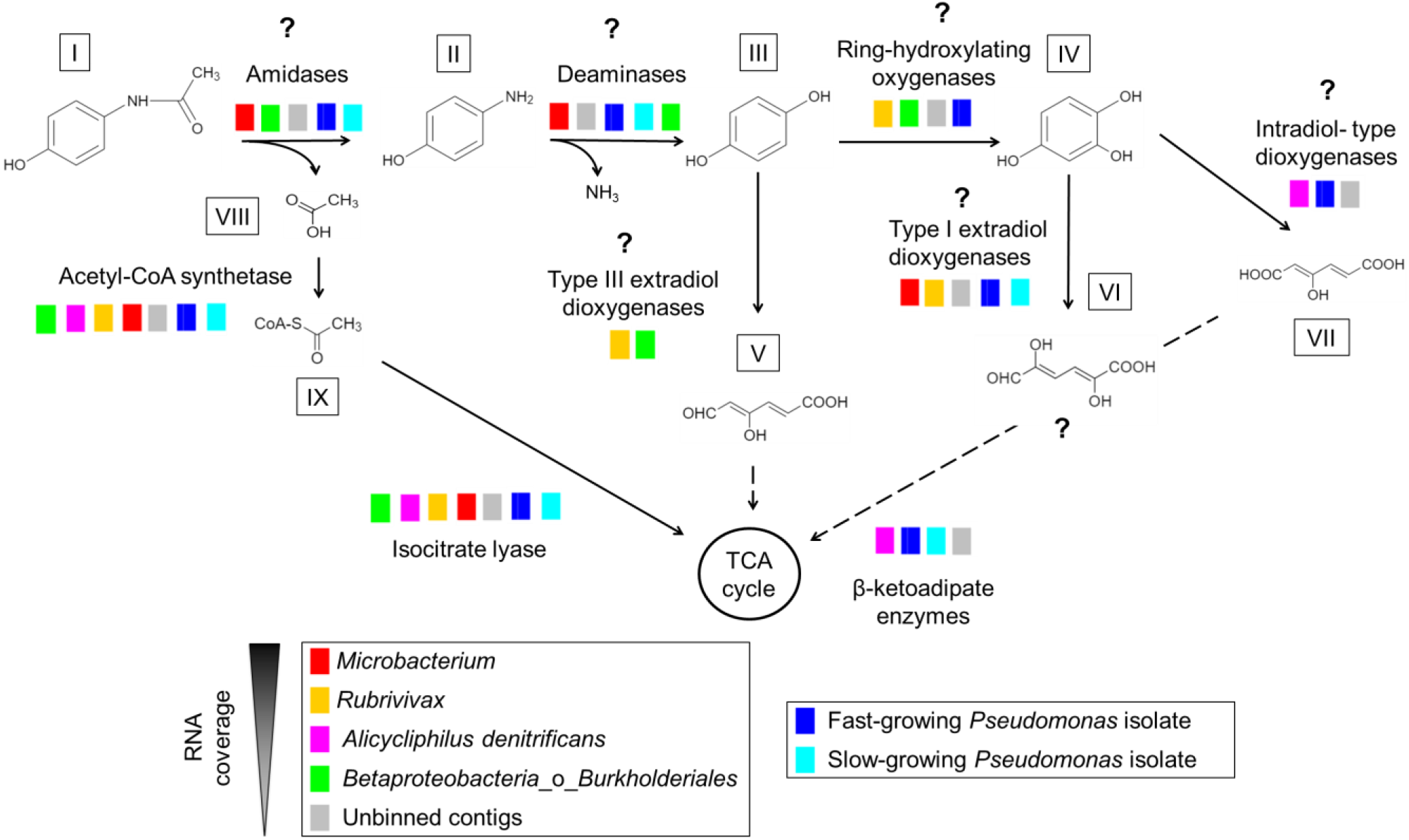
Paracetamol degradation pathway by the bioreactor microbial community and the Pseudomonas isolates. The question marks represent candidate enzymes and metabolites. Dashed lines correspond to conversions requiring more than one step. I paracetamol; II 4-aminophenol; III hydroquinone; IV hydroxyquinol or 1,2,4-trihydroxybenzene; V 4-hydroxymuconic semialdehyde or 4-hydroxy-6-oxo-2,4-hexadienoic acid; VI 2,5-dihydroxy-6-oxo-2,4-hexadienoic acid; VII 3-hydroxy-cis,cis-muconate or 3-hydroxy-2,4-hexadienedioic acid; VIII acetate; IX acetyl-CoA; TCA tricarboxylic acid.

### 3.7. Highly-expressed nitrification and denitrification genes in the bioreactor

The majority of the MAGs, except for *Patescibacteria* and *Myxococcales*_1, had highly expressed genes encoding enzymes from the denitrification pathway. All four genes encoding the full pathway of denitrification could only be found in one MAG, affiliated with *Alicycliphilus denitrificans*, as well as the in the unbinned contigs: nitrate reductase, nitrite reductase, nitric oxide reductase, and nitrous oxide reductase. The bioreactor was fully aerated, but biomass was spatially organized in small granules (Figure S3), so there might have been anoxic conditions towards the inside of the granules favoring denitrification. Nitrate and nitrite were not added to the medium, so nitrifying microorganisms were apparently also active in the bioreactor. A single highly expressed ammonia monooxygenase (subunits A, B, C; KGEGGEOM_12202, KGEGGEOM_12201, KGEGGEOM_12203) was encoded in the unbinned contigs, and affiliated with the complete ammonia oxidizer (comammox) *Nitrospira* sp. This finding suggests that some ammonia released from decaying biomass, from the paracetamol degradation and from the ammonia supply in the medium intended for assimilation (~1 mM) was converted into nitrate by comammox *Nitrospira* sp. and subsequently available for (oxygen-limited) denitrification.

## CONCLUSIONS

On the basis of our cultivation and metagenomic analysis, we conclude that APAP was immediately degraded by the activated sludge of a hospital WWTP and that a diverse microbial community was enriched under low APAP concentrations in a membrane bioreactor. High APAP concentrations in batch led to the dominance of a fast-growing *Pseudomonas* species. Several uncharacterized amidases from the AS family were highly expressed in the genome of a fast- and a slow-growing *Pseudomonas* species and the bioreactor metagenome. They might be cleaving APAP into 4-AP at different rates. Genes encoding for uncharacterized RidA family proteins were highly expressed in the genome of the *Pseudomonas* isolates and several bioreactor MAGs. They are known to have deaminase activity, so they might be converting 4-AP to HQ or cleaving reactive enamine intermediates. Genes encoding for intradiol- and extradiol-type dioxygenases were highly expressed in the genomes of the *Pseudomonas* isolates and the bioreactor metagenome. Many of these genes are part of the degradation pathway of aromatic amino acids. Therefore, microorganisms might take advantage of the side activities of existing enzymes encoded in their genomes for the degradation of APAP transformation products. Candidate APAP-degrading amidases, deaminases, and dioxygenases were not combined in the same gene cluster. Highly expressed genes encoding amidases were often found in the vicinity of mobile genetic elements, which suggests that APAP-degrading amidase genes are currently being exchanged between different bacteria via HGT.

Taken together, these results suggest a role of uncharacterized amidases, deaminases and dioxygenases in the biodegradation of APAP and the use of cross-feeding to efficiently degrade APAP in WWTP microbial communities. Furthermore, the high number of microorganisms able to degrade APAP might be the result of the broad substrate spectrum of amidases and its evolution, together with the fact that just one enzyme (amidase) is needed to grow on APAP-derived acetate. This study contributes to a better understanding of microbial evolution towards pharmaceutical biodegradation and demonstrates the complexity of this process due to the broad substrate spectrum of the involved enzymes.

## List of abbreviations used

WWTP: wastewater treatment plant
MBR: membrane bioreactor
GAC: granular activated carbon
APAP: N-acetyl-p-aminophenol or paracetamol
4-AP: 4-aminophenol
HQ: hydroquinone
HRT: hydraulic retention time
SRT: solid retention time
Pfast: *Pseudomonas* sp. isolate growing fast on APAP as sole carbon source
Pslow: *Pseudomonas* sp. isolate growing slow on APAP as sole carbon source.
HGT: horizontal gene transfer
MAG: metagenome-assembled genome
TPM: transcripts per million

## AUTHOR CONTRIBUTIONS

ARM, MJ, HOdC, and CW contributed to the conceptual framework of the manuscript. ARM conducted the experiments and data analysis. GS, GC, and HOdC contributed to bioinformatics analyses. TvA performed DNA and RNA Illumina sequencing of the slow-growing *Pseudomonas*. ARM wrote the manuscript with input from all the authors.

## DATA AVAILABILITY

All raw sequencing data (DNA and RNA) have been deposited at the read sequence archive (SRA) database of the NCBI under the BioProject ID PRJNA831879. The amino acid sequences and annotation of all genes in the metagenome and the *Pseudomonas* spp. genomes are deposited in Dans Easy (https://doi.org/10.17026/dans-xwd-fbj5). This dataset also contains the TPMs of the bioreactor and the *Pseudomonas* spp. transcriptomes and the amino acid sequences of the amidase genes used to create the phylogenetic trees in Figure 3 and supplementary Figure S2.

## ACKNOWLEDGEMENTS

The authors thank Stefan Hertel and Erwin Koetse for letting us take sludge from the MBR and GAC of Pharmafilter. We also thank Rob de Graaf for sharing his expertise with the HPLC and Guylaine Nuijten for her help with bioreactors. This research was supported by NWO-TTW grant 15759 and NWO/OCW grant SIAM 024002002.

