## Supplementary file 1 for "Microbial paracetamol degradation involves a high diversity of novel amidase enzyme candidates"

### Supplementary material of "Metagenomic and transcriptomic analysis of paracetamol biodegradation in a microbial community of a hospital wastewater treatment plant"

Ana B. Rios-Miguel[a], Garret J. Smith[a], Geert Cremers[a], Theo van Alen[a], Mike S.M. Jetten[a,b], Huub J. M. Op den Camp[a], Cornelia U. Welte[a.b]

[a]Department of Microbiology, Radboud University, Radboud Institute for Biological and Environmental Sciences, Heyendaalseweg 135, 6525 AJ Nijmegen, The Netherlands

[b]Soehngen Institute of Anaerobic Microbiology, Radboud University, Heyendaalseweg 135, 6525 AJ Nijmegen, The Netherlands

The supplementary material consists of Supplementary file 1 (this one including Figure S1, Figure S2, Figure S3 and Table S1) and Supplementary file 2 (amino acid sequence alignment of amidases)

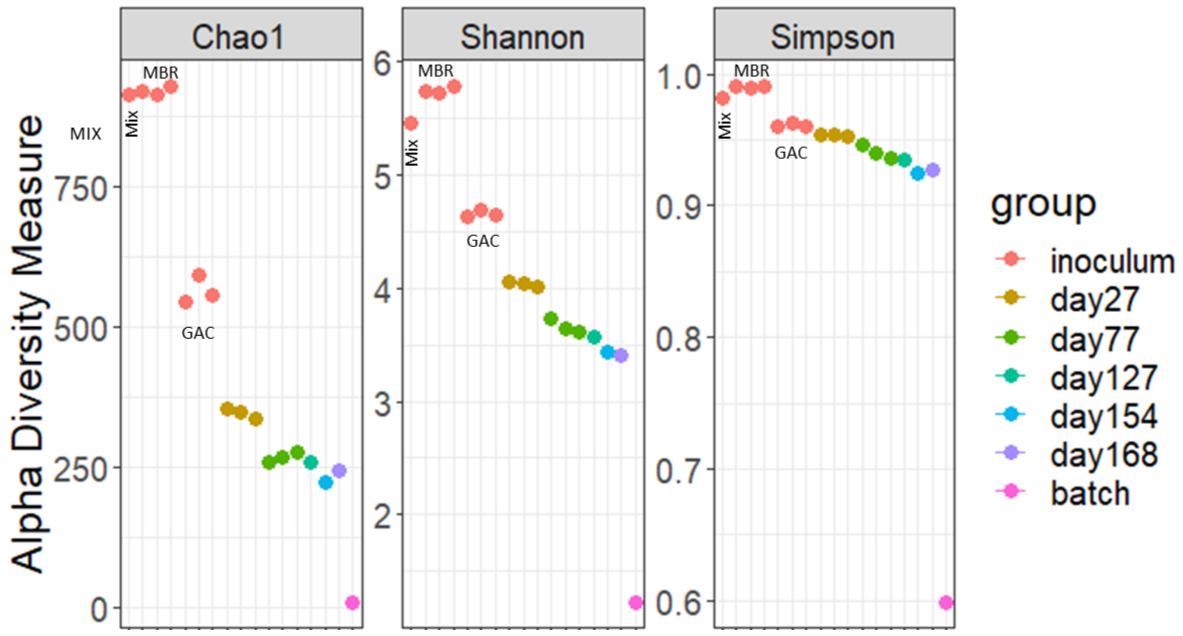


**Figure S1**. Alpha diversity of the inoculum (mix, membrane bioreactor (MBR), and granular activated carbon (GAC)), the bioreactor at different time points, and the biomass dilution in 400 mg/L of paracetamol (batch).

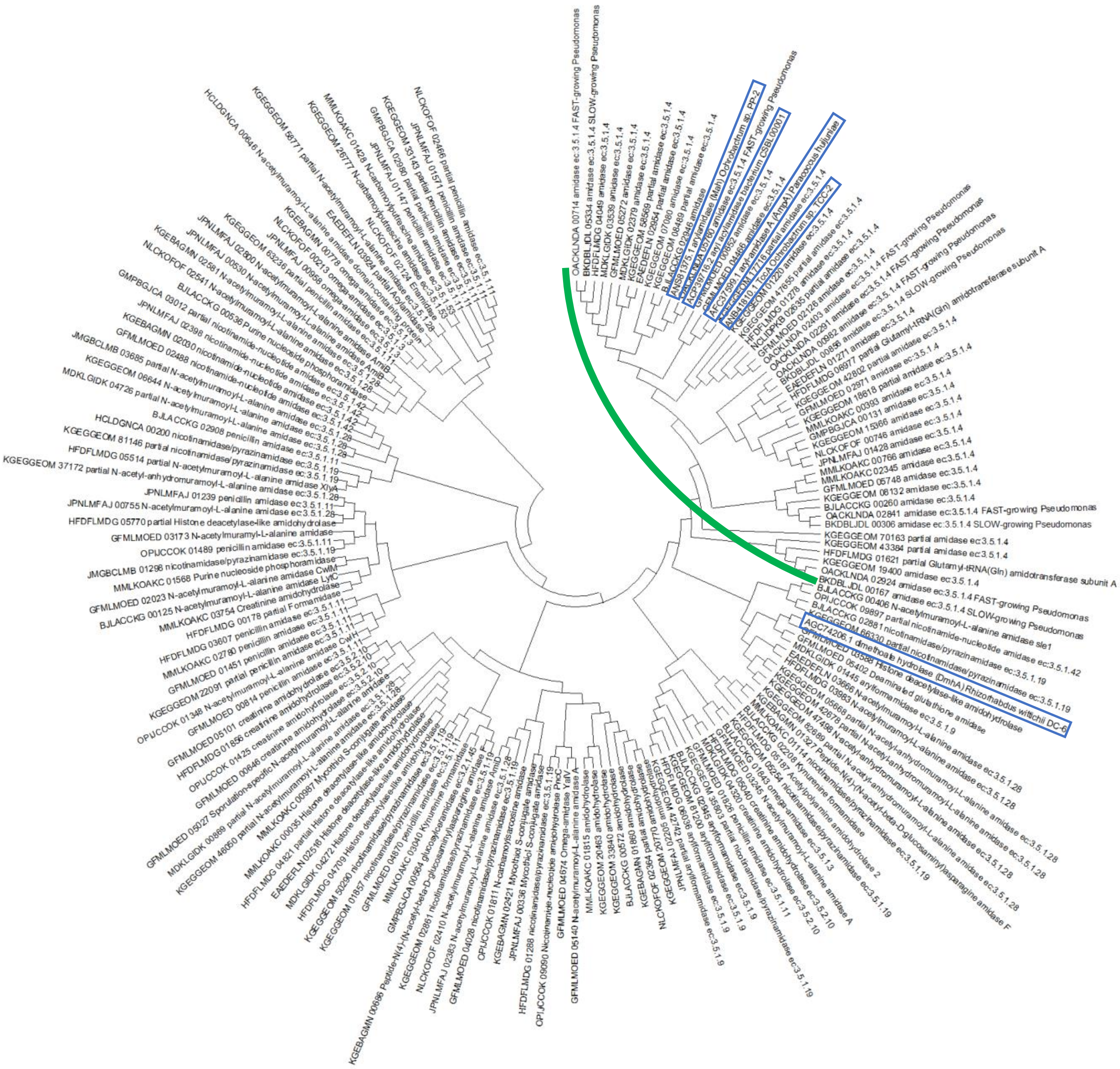

**Figure S2**. Phylogenetic tree of the top 150 most expressed amidases in the bioreactor, the uncharacterized amidases in the *Pseudomonas* isolates, and the amidases known to degrade paracetamol (in blue). The tree was created with the neighbor-joining method to analyze the topology (Saitou and Nei, 1987).

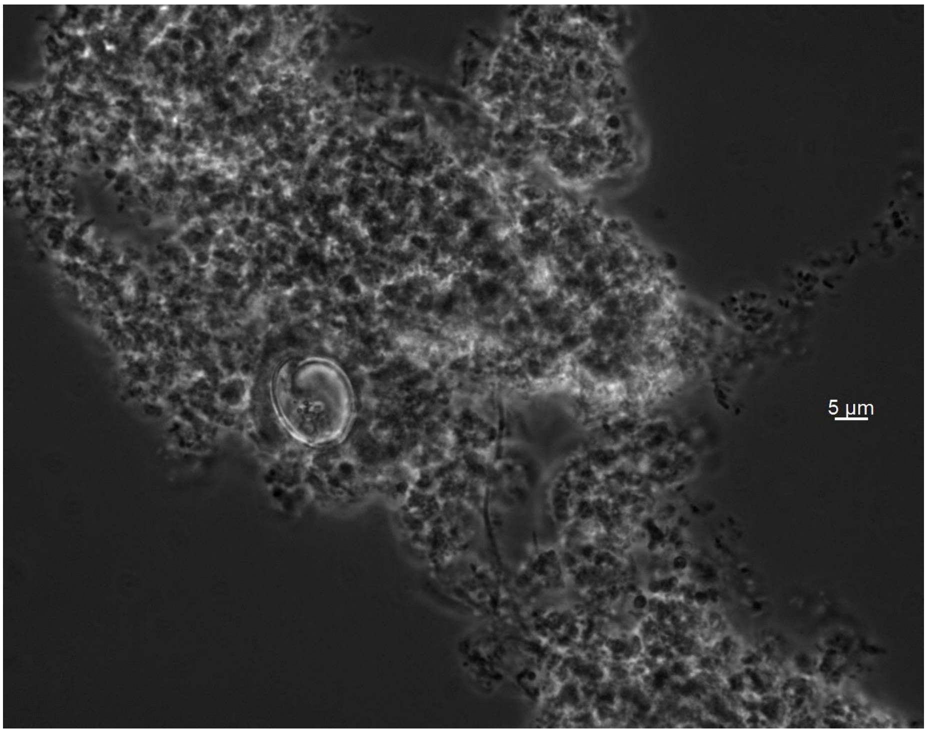


**Figure S3**. Granule of the bioreactor biomass.

**Table S1**. Amidase proteins known to degrade paracetamol (4-acetaminophenol). We did not take into account amidases able to degrade paracetamol but whose amino acid sequence was not reported.

| NCBI protein accession number | Microorganism | Isolation site | Substrates | Reference |
|---|---|---|---|---|
| ACP39716.2/ AYE89271.1, WP091572988.1 | Putative *Pseudomonas* sp. /*Comamonadaceae*. (*Proteobacteria*) | Soil in South Korea/ China | Paracetamol, 4-Nitroacetanilide, phenacetin, 4-Chloroacetanilide, acetanilide, Methyl N-(3,4-dichlorophenyl)carbamate (swep) | (Ko et al., 2010; Lee et al., 2015)/ (Zhang et al., 2020) |
| AFC37599.1 (AmpA) | *Paracoccus huijuniae* (*Alphaproteobacteria*) | Activated sludge from China | Paracetamol, propanil, dimethoate, omethoate, (chlor)propham, diflubenzuron, hexaflumuron, formamide and propionamide. NOT Carbofuran, carbaryl, diuron, linuron, metsulfuron-methyl, acetochlor and butachlor | (Zhang et al., 2012) |
| AGC74206.1 (DmhA) | *Rhizorhabdus wittichii* (*Alphaproteobacteria*) | Activated sludge from China | Paracetamol, propanil, dimethoate. NOT iflubenzuron, (chlor)propham, and linuron | (Chen et al., 2016) |
| ANB41810.1 (TccA) | *Ochrobactrum* sp. (*Alphaproteobacteria*) | River sediment from China | Paracetamol, triclocarban, diflubenzuron, (4,4’-dichloro)carbanilide, (chlor)propham, 4- | (Yun et al., 2017) |

| | | | | |
|---|---|---|---|---|
| | | | chlorophenylurea, 1-(3,4-dichlorophenyl)urea, 4-bromophenylurea, (4-chlor)acetanilide, forchlorfenuron. NOT barban, 4-fluorophenylurea, (2,6-dichloro)benzamide, propyzamide, chloramphenicol, florfenicol | |
| ANS81375.1 (Mah) | *Ochrobactrum* sp. (*Alphaproteobacteria*) | Soil from China | Paracetamol, propanil, (chlor)propham. NOT diuron and linuron | (Zhang et al., 2019) |


Chen, Q., Chen, K., Ni, H., Zhuang, W., Wang, H., Zhu, J., He, Q. and He, J. 2016. A novel amidohydrolase (DmhA) from Sphingomonas sp. that can hydrolyze the organophosphorus pesticide dimethoate to dimethoate carboxylic acid and methylamine. Biotechnology Letters 38(4), 703-710.

Ko, H.-J., Lee, E.W., Bang, W.-G., Lee, C.-K., Kim, K.H. and Choi, I.-G. 2010. Molecular characterization of a novel bacterial aryl acylamidase belonging to the amidase signature enzyme family. Molecules and Cells 29(5), 485-492.

Lee, S., Park, E.-H., Ko, H.-J., Bang, W.G., Kim, H.-Y., Kim, K.H. and Choi, I.-G. 2015. Crystal structure analysis of a bacterial aryl acylamidase belonging to the amidase signature enzyme family. Biochemical and Biophysical Research Communications 467(2), 268-274.

Saitou, N. and Nei, M. 1987. The neighbor-joining method: a new method for reconstructing phylogenetic trees. Mol Biol Evol 4(4), 406-425.

Yun, H., Liang, B., Qiu, J., Zhang, L., Zhao, Y., Jiang, J. and Wang, A. 2017. Functional Characterization of a Novel Amidase Involved in Biotransformation of Triclocarban and its Dehalogenated Congeners in Ochrobactrum sp. TCC-2. Environ Sci Technol 51(1), 291-300.

Zhang, J., Yin, J.-G., Hang, B.-J., Cai, S., He, J., Zhou, S.-G. and Li, S.-P. 2012. Cloning of a novel arylamidase gene from Paracoccus sp. strain FLN-7 that hydrolyzes amide pesticides. Appl Environ Microbiol 78(14), 4848-4855.

Zhang, L., Hang, P., Zhou, X., Dai, C., He, Z. and Jiang, J. 2020. Mineralization of the herbicide swep by a two-strain consortium and characterization of a new amidase for hydrolyzing swep. Microbial Cell Factories 19(1), 4.

Zhang, L., Hu, Q., Hang, P., Zhou, X. and Jiang, J. 2019. Characterization of an arylamidase from a newly isolated propanil-transforming strain of Ochrobactrum sp. PP-2. Ecotoxicology and Environmental Safety 167, 122-129.
