## Supplementary file 2 for "Microbial paracetamol degradation involves a high diversity of novel amidase enzyme candidates"

CLUSTAL O(1.2.4) multiple sequence alignment

Based on the crystal structure of ACP39716.2 in a soil bacterium, we marked the following regions:

- Canonical catalytic triad: Ser$^{187}$-*cis*Ser$^{163}$-Lys$^{84}$.
- Gly/Ser-rich motif: **GGSSGG**.
- The oxyanion hole: **[G]GGS**.
- Loop 1: Asp$^{85}$ – Ser$^{114}$.
- Loop 2: Pro$^{202}$ – Leu$^{222}$.
- *α-helix* (α10): Asp$^{354}$-Val$^{363}$
- Substrate-binding residues: Tyr$^{136}$ and Thr$^{330}$.

```
Unbinned_Comamonadaceae_KGEGGEOM_08132              ----MTNVNSDVIDLTELTVERVQAGFASGAFTSETLTKAYLKRIEEFNPSYNAIVFFN-      55
Paracoccus_huijuniae_AFC37599.1_AmpA                ---------MIPRLTNEPGAIEVAAQIRAGELSPLEAANAAIARIEALDGPLNAVVVRDF      51
Ochrobactrum_sp.TCC-2_ANB41810.1_TccA               --------MTATDELYYLPVTELSELIAERRLAPSELMSAVIARAEEVNPKLNALVAQRF      52
SLOW-growing_Pseudomonas_BKDBLJDL_05334             --------MTQKTAIGELTAVELLELYQRRQLSPVEVVDDVLARIDLHNPAVNAYCHVDG      52
Unbinned_Actinomycetia_KGEGGEOM_07080               ---------MDPSEYAWMSATDLVKALQSKQLSPVDLIDAAIARIEQRNPAINAVVHLGA      51
Unbinned_Actinomycetia_KGEGGEOM_08489_partial       ------VIDHRNLPNRWVSATETVRRITSGEQSATEVTTDSIHRIEHLNSMYNAVTFSDP      54
Microbacterium_BJLACCKG_02946                       VNADPVSIASRHTAGAANTATETARRVRAGEISALEVTEEALARIDQVNPALGAVTFVAA      60
Ochrobactrum_sp.PP-2_ANS81375.1_Mah                 ---------MERSDLDYASATEIARLVRTRQISAADVTEHAISRIEARNGSLNAFVYTDF      51
FAST-growing_Pseudomonas_OACKLNDA_05760             ---------MIGSQVHWKTATEIAGLVKSKQISPREVVEGTIELIEQRNPSMNAVTYKAY      51
Crystal_structure_bacterium_CSBL00001_ACP39716.2    -------MGKSHSPVHWKSAAEIVELVKSKQISPREVVESTIDLIEQRDPGLNAVVYKAY      53
                                                                       .            :        :   :  :   .*

Unbinned_Comamonadaceae_KGEGGEOM_08132              EKAVEEARAIDARRAAGEKLGPLAGVPVVVKEAMD-MKGFPTTGGWSLLYSKTGGVDLLP      114
Paracoccus_huijuniae_AFC37599.1_AmpA                DRARDAARELDGQPA---EDRPLFGVPMTVKESFD-VAGLPTTWGHVPFKDY---RPT--      102
Ochrobactrum_sp.TCC-2_ANB41810.1_TccA               EAATREAAAADN----EPSRGVLHGIPITLKDLAYETPDLPSTYGSRAFAGY---EP---      102
SLOW-growing_Pseudomonas_BKDBLJDL_05334             EGARAAAARASEQRWQRGQPCGRLDGVPASIKDLTL-TRGMPTRKGSRTTS-----GSGPW      106
Unbinned_Actinomycetia_KGEGGEOM_07080               EDARAAARTAERALA-EGRAGPLAGLPVLMKDLFDFKPGWPSTLGGVPALAD---NIAQ-      106
Unbinned_Actinomycetia_KGEGGEOM_08489_partial       DLALSTARDIDRRLAASDEVGPLAGVPTLMKDLYGFRPGWPTTLGGLSAARG---DTASE      111
Microbacterium_BJLACCKG_02946                       DQARDAAARLDRRIRSGESVGPLAGVPTLVKDLYGFVPGWPSTLGGIEALRD---HRTPD      117
Ochrobactrum_sp.PP-2_ANS81375.1_Mah                 EQARSRAKDLDTRISAGEDVGPLAGVPTAIKDLFNFYPGWPSTLGGIRCLRD---FKL--      106
FAST-growing_Pseudomonas_OACKLNDA_05760             DEARVKAAELERHIMNGGRTGALAGVPTLMKDLFAAKPGWPSTLGGISALKH---LRGAE      108
Crystal_structure_bacterium_CSBL00001_ACP39716.2    DEAREKAAALERRIMQGEPVGMLAGVPTLMKDLFAAKPGWPSTLGGIRALKD---ARGAA      110
                                                    : *   *   :           * *:*  :*:      . *:  *
```

```
Unbinned_Comamonadaceae_KGEGGEOM_08132          ETDSPVVARMRQADTVILGKTNIPILSHTGAHANGSWAGSTYNSAGREFLPGGSSAGTAT    174
Paracoccus_huijuniae_AFC37599.1_AmpA            -RDARVVQLLKDAGAIILGKTNVPPDLADMQSNNP-VYGRTNNPYDHSRVAGGSSGGSAV    160
Ochrobactrum_sp.TCC-2_ANB41810.1_TccA           GFETVVGQRLRAAGTIAIGRTNSPEFGLTNTCESA-QFGPTSNPWRPEHTPGGSSGGAGA    161
SLOW-growing_Pseudomonas_BKDBLJDL_05334         EIDAPFSAFMREAGAVLVGKTTTPEFGWKGVTDNP-LYGITRNPWDTRLTAGGSSGGAAA    165
Unbinned_Actinomycetia_KGEGGEOM_07080           -HSCLFVERMESAGAIVLGKTNSPVFGARGITDNP-LFGPTCNPFDVSRNPGGSSGGAAA    164
Unbinned_Actinomycetia_KGEGGEOM_08489_partial   GAWSRFPKQVTADDGVLLGQTNSSTFGFSGVTDNA-LFGPSRNPFEPTRNTGGSSGGSAA    170
Microbacterium_BJLACCKG_02946                   GLWSAYPRSAVSADAILIGQSNSPVFGFRGVTDNT-LFGPTRNPFDLTRNAGGSSGGAAA    176
Ochrobactrum_sp.PP-2_ANS81375.1_Mah             DVKSRYATKMEEAGAVVLGITNSPVLGFRGTTDND-LYGPTRNPFDLSRNSGGSSGGTSA    165
FAST-growing_Pseudomonas_OACKLNDA_05760         GAWSTFPLKMSNEDSLLLGQTNSPVFGFRGVTDNK-FFGPTRNPFNLDYNAGGSSGGAAA    167
Crystal_structure_bacterium_CSBL00001_ACP39716.2 GVWSTYPLKMSGEDSLLLGQTNSPVYGFRGTTDNT-FFGPTRNPFNLDFNAGGSSGGAAA    169
                                                             . : :* :.           .    * : *        ****.*:..

Unbinned_Comamonadaceae_KGEGGEOM_08132          AVGANFCVLGLAEETAGSIQNPASAQGLVGIKPTFGLVPNAGVMP--------LASLRDV    226
Paracoccus_huijuniae_AFC37599.1_AmpA            AVATGMVPAEYGSDIGSSIRNPAHFNGIYGHKTTFGLVSRRGHGHPVAGGKDMHAGPLSV    220
Ochrobactrum_sp.TCC-2_ANB41810.1_TccA           AVAAGIAPLAAANDGGGSCRVPASSCGVVGLKPSRGRVPWAPTSY-----EYW--AGFAT    214
SLOW-growing_Pseudomonas_BKDBLJDL_05334         AAALNLGVLHQGSDAGGSIRIPCAFTGTFGIKPTFGYVPQWPASA-----MTV----LSH    216
Unbinned_Actinomycetia_KGEGGEOM_07080           AVAAGFVPIAEGTDAGGSIRIPAAWTSTYGFKPSAGRVPSIIRPL-----GFMTAAPFIT    219
Unbinned_Actinomycetia_KGEGGEOM_08489_partial   AVASGMIPVAGASDAGGSIRIPAAWTNTVGFQPSAGRVPSTPRPA-----LFHL-GPHLY    224
Microbacterium_BJLACCKG_02946                   AVATGIVPVAGASDAGGSIRIPAAWTNTVGFQPSAGRVPSTPRPV-----GFHL-APFLY    230
Ochrobactrum_sp.PP-2_ANS81375.1_Mah             AVADGLLPIGDGTDGGGSIRIPAAWCHVFGFQASPGRIPLAIRPN-----AFGAAAPFIY    220
FAST-growing_Pseudomonas_OACKLNDA_05760         VVADGIVPIAGGTDAGGSVRIPAAWTNTYGFQPSIGRIPFISRPN-----AFHL-ATYIY    221
Crystal_structure_bacterium_CSBL00001_ACP39716.2 LVADGIVPVAGGTDGGGSIRIPAAWTNTYGFQPSIGRVPFKSRPN-----AFHP-GPYLY    223
                                                 .. .:     . : ..* : *.      * : : * :

Unbinned_Comamonadaceae_KGEGGEOM_08132          VGPIARCVRDAALTLDVLAGFSMEDPKTNASVGRRPKGGYASKLDKGALAGKRIGLYGP-    285
Paracoccus_huijuniae_AFC37599.1_AmpA            TGPLARSAEDLQLLLQVTAERP-------------------LARRRKSLTEMRFLAVLD-    260
Ochrobactrum_sp.TCC-2_ANB41810.1_TccA           NGPIARTVEDVALLLDAMSGPVVGEPYGLP-----APSESFLTASRRRPGPLRIAFSCTP    269
SLOW-growing_Pseudomonas_BKDBLJDL_05334         LGPMTRTVDDSVLMLDCVARPDARDGLAGA-----PRQAPWLSQQ-QDLSGLRIAYSAN-    269
Unbinned_Actinomycetia_KGEGGEOM_07080           EGPITRTVADAALAMTALAGPDSRAPHCLD-----GVLDYRAALD-GSIAGRRLGYSPN-    272
Unbinned_Actinomycetia_KGEGGEOM_08489_partial   EGPITRTVEDAVLLMNSLQGYDPRDPYAAP-----APAFSPALLS-SGVRGKRIGLSLD-    277
Microbacterium_BJLACCKG_02946                   EGPITRTVQDAALVIDALSSHDPHDPTSVD-----GLPSMSEAIT-RDIDELRIGIVED-    283
Ochrobactrum_sp.PP-2_ANS81375.1_Mah             EGPITRTVEDAALAMSVLAGSDPADPFSLN-----DRLDWLGAVD-QPITSLRIGFTPD-    273
FAST-growing_Pseudomonas_OACKLNDA_05760         EGPITRTVEDAALAMNALHGFDRRDPNSLR-----VKLDFTSALV-QGVRGKKIGLTLD-    274
Crystal_structure_bacterium_CSBL00001_ACP39716.2 EGPITRTVRDAALAMNVLHGFDRRDPASLR-----VKLDFTSALA-QGVRGKKIGLTLN-    276
                                                 **::* . *  * :                                     ::
```

```
Unbinned_Comamonadaceae_KGEGGEOM_08132               GWRNSTFGEETISLYARAQEELKKLGATFVNDPFAGSGLRELRKPIMPGAEFDARGMESI      345
Paracoccus_huijuniae_AFC37599.1_AmpA                 -HPSSAIDASVRGPIEAALAEIERAGASVDRAS-------------------------ALL      295
Ochrobactrum_sp.TCC-2_ANB41810.1_TccA                PKPHDRLNAEVKQTFLAAVANFEALGHTVTEID-------------------------HGL      305
SLOW-growing_Pseudomonas_BKDBLJDL_05334              -FGYVQVAPQIQALVAQAVQRLARLGAQVEEVD-------------------------PGF      304
Unbinned_Actinomycetia_KGEGGEOM_07080                -LGAFPVEPAVARAVEHALTGFEECGAQVQQTE-------------------------VSL      307
Unbinned_Actinomycetia-KGEGGEOM_08489_partial        -FGGFPVNPLVRDTITEAAATFEQLGAIVDPID-------------------------VSL      312
Microbacterium_BJLACCKG_02946                        -FGGFPVDPAVRATVRRAADAFSGVAKSVSAVS-------------------------LNL      318
Ochrobactrum_sp.PP-2_ANS81375.1_Mah                  -FGGFPVEPAVAATIAHAVRAFEQAGAKIVPLK-------------------------LDF      308
FAST-growing_Pseudomonas_OACKLNDA_05760              -YGVFPVQPEIKDLISKTAQVFTQLGAHVEFVD-------------------------LGI      309
Crystal_structure_bacterium_CSBL00001_ACP39716.2     -YGVFPVQQEIQDLIGKAARVFTELGAHVEFVD-------------------------LGI      311
                                                           .          :   :   .  .                              :

Unbinned_Comamonadaceae_KGEGGEOM_08132               PYDIEKYLQRMGANVALKTFADFAKATEKEGAFGPNGVLSYMPHSPEFV-EVMKN-P---      400
Paracoccus_huijuniae_AFC37599.1_AmpA                 PDLAEQHYN-YMRLVNVAMMRGNPG------PLAEQHANYMR----LVNVAMMRGNPGPA      344
Ochrobactrum_sp.TCC-2_ANB41810.1_TccA                DGIFDSFIR----VIAANTALSVTQ------TVPLGSLNLLE--PNTLGLAQRGW-G---      349
SLOW-growing_Pseudomonas_BKDBLJDL_05334              SDPLETFNTLWFAGAARLASAL-------------------SDEQKALLDPGLRW-IAEQ      344
Unbinned_Actinomycetia_KGEGGEOM_07080                PADHEELAALWHRSMMQVTLASL-E------GMQAQGIDLLADLAAHFPPALAEQ-IEQ-      358
Unbinned_Actinomycetia_KGEGGEOM_08489_partial        TYTHDELTQMWLRSMGLLMLADL-D------SLRARGRAL-A--PRDLPEPVLYW-TEV-      360
Microbacterium_BJLACCKG_02946                        GQSHDELTETWLRMMGTAMLSEM-D------SHRRQKVDLTA--AGGVPLEVLRW-TSR-      367
Ochrobactrum_sp.PP-2_ANS81375.1_Mah                  GYTHDELSQLWCRMISQGTIAVV-D------SFAENGLHL----EPDFPAPVMEW-AQK-      355
FAST-growing_Pseudomonas_OACKLNDA_05760              PYSQEQMSDAWCRMLAIPTAASM-H------ALHEDGIDLFSEHRADIPDALMKW-IDA-      360
Crystal_structure_bacterium_CSBL00001_ACP39716.2     PYSQKQMSDAWCRMIAIPTVASM-Q------ALRKEGIDLYGEHRADIPDALMKW-IDA-      362
                                                         .                                          .

Unbinned_Comamonadaceae_KGEGGEOM_08132               ---SNPPD-MASFIALRETYLDIFNSVFDKHKLDAV----IYPQMRGPLGPLHGDEV---      449
Paracoccus_huijuniae_AFC37599.1_AmpA                 GQTMALADYYQLLDTQE-CNRYAWADLFAEYDFVLAPPLPFVAYPHDATPIYERRIPING      403
Ochrobactrum_sp.TCC-2_ANB41810.1_TccA                ---LSAMDYCEAINHLRTTAALSMARWTEDFDVLLTPTLTDLPPLTGQMPSYDGDL---D      403
SLOW-growing_Pseudomonas_BKDBLJDL_05334              GAQISLGE-YTQALEARAELIAKMNAFHQRYDVLVSPMLPL---VAFEAGHNVP----PG      396
Unbinned_Actinomycetia_KGEGGEOM_07080                TRTMTARQ-VRIDDVLRTEIYETLTSAINSFDLLLTPTVGGLPVVNASRGETIGPTQLNG      417
Unbinned_Actinomycetia_KGEGGEOM_08489_partial        AAAMSAAD-VRSDRVMRTAVLDGLVDAMENHDLLVGPTVVDLPVLNSSDELTIGPTTVDG      419
Microbacterium_BJLACCKG_02946                        AMAMTMRE-LLADRETRTAVLDGFLSAMDGVDLLVGPTVTALPVLNSTGGRTVGPSEVDG      426
Ochrobactrum_sp.PP-2_ANS81375.1_Mah                  AKNATPLD-LHRDQVMRTKVYDVLNAAFSQVDLIAGPTTTCLPTPNGERGMTVGPSEIAG      414
FAST-growing_Pseudomonas_OACKLNDA_05760              VADINVQQ-ISADQILRTSVFDCMNRVFDRFDLLLAPTLACMPVRNATDGSTEGPSAING      419
Crystal_structure_bacterium_CSBL00001_ACP39716.2     VADISVQQ-ISADQLLRTTVFDCMNGVFDRFDLLLAPTLACMPVRNATDGCTEGPSQING      421
                                                            :        .              ..
```

```
Unbinned_Comamonadaceae_KGEGGEOM_08132               ----IDA-----MVVSEINIAGLPAVTVPAGFYASGAPFNLVIVAPQWSEADILALAYAY      500
Paracoccus_huijuniae_AFC37599.1_AmpA                 KDSAFADALAWAG---LANFPNLPSTVVPVGES-AGLPCGMQVMGPEWSDLDCIAAAGAI      459
Ochrobactrum_sp.TCC-2_ANB41810.1_TccA                --ACYLHMLGHNAFTYPFNVTGQPALSIPCGWSTSGLPIGLQIIGGMGQEARVLALAAAY      461
SLOW-growing_Pseudomonas_BKDBLJDL_05334              --SGMAQWMEWTPFSYPFNLTQQPAASVPCGFTREGLPVGLQVVAGRFADEQVLRVCKVY      454
Unbinned_Actinomycetia_KGEGGEOM_07080                --QAVDPMIGWAL-TFPFNFTGHPAASVPAGLV-DGLPVGMQLIGRQMGDLDVLTFSAAY      473
Unbinned_Actinomycetia_KGEGGEOM_08489_partial        --TAVNPLIGWCP-TYLTNFSGSPSVSVPAGFA-EGLPVGMLIIGRKHRDEDVLAAAAAL      475
Microbacterium_BJLACCKG_02946                        --VPVDPLIGWCP-TYLTNFTGMPSISLPAGFA-ENLPVGLLIIGRKYRDAEVFTAAAAF      482
Ochrobactrum_sp.PP-2_ANS81375.1_Mah                  --TPINRLIGFCP-TFLTNFTGNPAASLPAGLA-DGLPVGLMLIGPRRDDLTVLSASAAF      470
FAST-growing_Pseudomonas_OACKLNDA_05760              --EKVNPLIGWCM-TYLTNFSGHPSASVPAGLI-DGLPVGMLIIGDRQADLDVIAASAAF      475
Crystal_structure_bacterium_CSBL00001_ACP39716.2     --EEIDPLIGWCM-TYLTNFSGHPSASVPAGLI-DGLPAGMLIIGDRQADLDVIAASAAF      477
                                                                       *.   *:  :* *    . * .: ::.    :   :  . .

Unbinned_Comamonadaceae_KGEGGEOM_08132               EQGTTHRKAPELKKA-------- 515
Paracoccus_huijuniae_AFC37599.1_AmpA                 GALMEG----------------- 465
Ochrobactrum_sp.TCC-2_ANB41810.1_TccA                EEAHPWAARKPPL---------- 474
SLOW-growing_Pseudomonas_BKDBLJDL_05334              EQHYPSRHLQAPITG-------- 469
Unbinned_Actinomycetia_KGEGGEOM_07080                ERARPWVGNYAQVDG-------- 488
Unbinned_Actinomycetia_KGEGGEOM_08489_partial        ETAR------------------- 479
Microbacterium_BJLACCKG_02946                        EKIRPWHGAYREIAIGNASVQPD 505
Ochrobactrum_sp.PP-2_ANS81375.1_Mah                  ERVQPWADSYRIPAARPLGSQ-- 491
FAST-growing_Pseudomonas_OACKLNDA_05760              EEARPWLQYYDIPARRALQ---- 494
Crystal_structure_bacterium_CSBL00001_ACP39716.2     ERASPWSQYYDIPAGRPL----- 495
```
